# Retroposon Insertions within a Multispecies Coalescent Framework Suggest that Ratite Phylogeny is <u>not</u> in the ‘Anomaly Zone’

**DOI:** 10.1101/643296

**Authors:** Mark S. Springer, John Gatesy

**Affiliations:** Department of Evolution, Ecology, and Organismal Biology, University of California, Riverside, CA 92521, USA; Division of Vertebrate Zoology, American Museum of Natural History, New York, NY 10024, USA; Sackler Institute for Comparative Genomics, American Museum of Natural History, New York, NY 10024, USA

**Keywords:** ASTRAL, retroposon, coalescence; gene tree, incomplete lineage sorting, species tree

## Abstract

Summary coalescence methods were developed to address the negative impacts of incomplete lineage sorting on species tree estimation with concatenation. Coalescence methods are statistically consistent if certain requirements are met including no intralocus recombination, neutral evolution, and no gene tree reconstruction error. However, the assumption of no intralocus recombination may not hold for many DNA sequence data sets, and neutral evolution is not the rule for genetic markers that are commonly employed in phylogenomic coalescence analyses. Most importantly, the assumption of no gene tree reconstruction error is routinely violated, especially for rapid radiations that are deep in the Tree of Life. With the sequencing of complete genomes and novel pipelines, phylogenetic analysis of retroposon insertions has emerged as a valuable alternative to sequence-based phylogenetic analysis. Retroposon insertions avoid or reduce several problems that beset analysis of sequence data with summary coalescence methods: 1) intralocus recombination is avoided because retroposon insertions are singular evolutionary events, 2) neutral evolution is approximated in many cases, and 3) gene tree reconstruction errors are rare because retroposons have low rates of homoplasy. However, the analysis of retroposons within a multispecies coalescent framework has not been realized. Here, we propose a simple workaround in which a retroposon insertion matrix is first transformed into a series of incompletely resolved gene trees. Next, the program ASTRAL is used to estimate a species tree in the statistically consistent framework of the multispecies coalescent. The inferred species tree includes support scores at all nodes and internal branch lengths in coalescent units. As a test case, we analyzed a retroposon dataset for palaeognath birds (ratites and tinamous) with ASTRAL and compared the resulting species tree to an MP-EST species tree for the same clade derived from thousands of sequence-based gene trees. The MP-EST species tree suggests an empirical case of the ‘anomaly zone’ with three very short internal branches at the base of Palaeognathae, and as predicted for anomaly zone conditions, the MP-EST species tree differs from the most common gene tree. Although identical in topology to the MP-EST tree, the ASTRAL species tree based on retroposons shows branch lengths that are much longer and incompatible with anomaly zone conditions. Simulation of gene trees from the retroposon-based species tree reveals that the most common gene tree matches the species tree. We contend that the wide discrepancies in branch lengths between sequence-based and retroposon-based species trees are explained by the greater accuracy of retroposon gene trees (bipartitions) relative to sequence-based gene trees. Coalescence analysis of retroposon data provides a promising alternative to the status quo by reducing gene tree reconstruction error that can have large impacts on both branch length estimates and evolutionary interpretations.

---

In a landmark paper on gene trees and species trees, Maddison (1997) likened phylogeny to a cloud of gene histories, a “cloudogram”, where individual gene trees may disagree with the containing species tree. Biological explanations for this discord include incomplete lineage sorting (ILS), hybridization (reticulation), and gene duplication followed by lineage extinction. Subsequent to Maddison (1997), numerous methods have been developed to reconstruct species trees with sets of heterogeneous gene trees. The majority of these methods are based on the multispecies coalescent and attempt to account for ILS (e.g., Liu et al., 2009a,b; 2010; Heled and Drummond, 2010; Mirarab et al., 2014c).

The development of ILS-aware approaches is important because the sorting of ancestral polymorphism may mislead concatenation (= supermatrix) methods, especially in the ‘anomaly zone’, a region of tree space where successive short internal branches (in coalescent units = CUs) permit the passage of alleles across multiple speciation events. In such circumstances, the most probable gene tree does not match the species tree and maximum likelihood (ML) supermatrix analysis fails with the accumulation of more and more data, while coalescence methods are statistically consistent (Degnan and Rosenberg, 2006, 2009; Mendes and Hahn, 2017). However, the simulations and mathematical work that demonstrate the statistical consistency of coalescence methods are based on several simplifying assumptions that likely do not hold for most empirical systematic datasets. If these assumptions are violated, then the perceived strengths of coalescence methods relative to concatenation may be offset by their own unique problems, and statistical consistency is not guaranteed (Gatesy and Springer, 2013, 2014; Patel et al., 2013; Mirarab et al., 2014a; Springer and Gatesy, 2014, 2016, 2018a, b; Bayzid et al., 2015; Mirarab and Warnow, 2015; Simmons and Gatesy, 2015; Gatesy et al., 2017, 2018; Scornavacca and Galtier, 2017; Molloy and Warnow, 2018).

Among coalescent methods, the application of full likelihood or Bayesian approaches such as *BEAST (Heled and Drumond, 2010) is challenging for genome-scale data sets because of a high computational burden (Knowles, 2009; Bayzid and Warnow, 2013; Zimmermann et al., 2014). Hence, summary coalescence methods that rely on estimated gene trees have emerged as popular alternatives (Liu et al., 2009a, 2010; Mirarab et al., 2014b). Widely used summary coalescence methods include STAR (Liu et al., 2009b), MP-EST (Liu et al., 2010), NJst (Liu and Yu, 2011), ASTRAL (Mirarab et al., 2014c; Mirarab and Warnow, 2015; Zhang et al., 2017), and ASTRID (Vachaspati and Warnow, 2015). These methods explicitly address the problem of ILS, but are not guaranteed to recover the correct species tree unless several model assumptions are true (Liu et al., 2009a; Roch and Warnow, 2015; Springer and Gatesy, 2018a).

Importantly, coalescence genes (c-genes), segments of the genome for which there has been no recombination over the phylogenetic history of a clade, are the fundamental units of analysis for summary coalescence methods (Doyle, 1992, 1997; Springer and Gatesy, 2016, 2018a). A foundational assumption of coalescence methods is that recombination occurs freely among c-genes but not at all within c-genes (Liu et al., 2009a). Thus, it is inappropriate to combine multiple c-genes in summary coalescence analyses, as for example when protein-coding sequences are stitched together from far-flung exons that in some cases are more than a megabase apart (Gatesy and Springer, 2013, 2014; Springer and Gatesy, 2016, 2018a; Scornavacca and Galtier, 2017). In addition, the length of c-genes becomes shorter with denser taxon sampling owing to the ‘recombination ratchet’ (Gatesy and Springer, 2014; Springer and Gatesy, 2016, 2018a). An important consequence of shrinking c-gene size with increased taxon sampling is that gene tree reconstruction accuracy decreases because there are increasingly fewer informative sites to resolve relationships among ever more taxa. However, the statistical consistency of summary coalescence methods is only guaranteed when gene trees are reconstructed accurately (Liu et al., 2009a,b, 2010; Roch and Warnow, 2015).

In analyses of DNA sequences, there also are non-biological reasons for artificially increased conflicts among gene trees. Some ML programs for tree inference (e.g., PhyML, RAxML) will return fully bifurcating gene trees even when there are no informative sites in a matrix (Simmons and Norton, 2013). Furthermore, long branch misplacement, sequence misalignments, orthology problems, tracts of missing data, insufficient informative variation, poor tree searches, and model mis-specification hinder the accurate reconstruction of gene trees (Huang and Knowles, 2009; Huang et al., 2010; Gatesy and Springer, 2014; Simmons and Gatesy, 2015; Hosner et al., 2016; Springer and Gatesy, 2016, 2018b,c; Gatesy et al., 2017; Sayyari et al., 2017). These diverse sources of gene tree heterogeneity lead to violations of another important assumption of summary coalescence methods, which is that ILS is the sole cause of topological conflicts among gene trees (Liu et al., 2009a). Indeed, ILS may be a minor contributor to gene tree heterogeneity for some empirical data sets (Scornavacca and Galtier, 2017).

Summary coalescent methods further assume that c-genes evolve neutrally, but this assumption is generally false for protein-coding sequences and ultraconserved elements (UCEs). UCEs are typically under extreme purifying selection (Katzman et al., 2007). Protein-coding genes, in turn, generally evolve under purifying, positive, or balancing selection (Bustamante et al., 2005; Piertney and Oliver, 2006). Nevertheless, UCEs and protein-coding genes are routinely analyzed in phylogenomic coalescence analyses using methods that assume neutral evolution (e.g., McCormack et al., 2012; Song et al., 2012; Liu et al., 2017).

Retroposon insertions provide an important alternative to sequence-based analyses for reconstructing species trees, and have been widely employed to investigate difficult phylogenetic problems within Mammalia (Nikaido et al., 1999, 2001; Nishihara et al., 2006, 2009; Churakov et al., 2010; Nilsson et al., 2012; Hartig et al., 2013; Doronina et al., 2015, 2017a, 2017b, 2019; Gallus et al., 2015; Lammers et al., 2019). The same approach increasingly has been applied to additional taxonomic groups including birds (Suh et al., 2011, 2015b; Matzke et al., 2012; Sackton et al., 2019) and crocodilians (Suh et al., 2015a). Although retroposon insertions have been analyzed using variants of parsimony (e.g., Nikaido et al., 1999; Suh et al., 2015b), distance methods (e.g., Lammers et al., 2019), or networks (e.g., Doronina et al, 2017a), they have several properties that are well matched to the assumptions of summary coalescence methods, with the caveat that these methods must permit polytomies in gene trees, because each retroposon insertion only provides information for a single bipartition of taxa. First, the presence/absence property of an individual retroposon insertion represents a single evolutionary event and is not subject to intralocus recombination. Second, gene tree reconstruction error is greatly reduced relative to standard sequence-based gene trees because retroposon insertions are virtually free of homoplasy (Shedlock and Okada, 2000; Shedlock et al., 2004; Hallström et al., 2011; Doronina et al., 2017b, 2019). Retroposon insertion at a specific genomic location is a rare event, as is the precise excision of an inserted sequence (Shedlock and Okada, 2000; Doronina et al, 2019). Third, retroposon insertions almost always occur in non-coding regions of the genome rather than in protein-coding regions that are under strong selection (negative or positive). Thus, many retroposon insertions may be effectively neutral (Kuritzin et al., 2016; Kubiak and Makalowska, 2017; Doronina et al., 2019). There are important exceptions, for example when retroposons provide new promoter sequences for a nearby gene (Dekel et al., 2015), but in the main, these elements occur in genomic ‘safe havens’ where they are tolerated (Chuong et al., 2017).

Here, we outline a general approach for analyzing retroposon insertion data that is fully consistent with the multispecies coalescent (Fig. 1) using data presented by Cloutier et al. (2019) and Sackton et al. (2019) as a test case. Their retroposon dataset for palaeognath birds (tinamous and ratites) is comprised of 4301 retroposon insertions that were scored for 12 ingroup species and a single outgroup (chicken). These inferred insertions provide support for 20 different bipartitions of taxa with counts that range from one to 2318. Retroposon distributions for Palaeognathae were taken from Cloutier et al.’s (2019) online document (“CR1_insertion_pattern_summary.xls”) to construct a presence/absence matrix for the 13 taxa. For each of the 4301 retroposon insertions, we created a gene tree with just a single split or bipartition (Fig. 1). We then executed analysis of the 4301 incompletely resolved gene trees using the summary coalescent method ASTRAL-III v. 5.6.1 (Zhang et al., 2017). This version of ASTRAL outputs the species tree, estimated branch lengths in CUs, and local posterior probabilities (Sayyari and Mirarab, 2016) (Fig. 1). As an additional measure of clade support, we employed the ‘gene-wise’ resampling option in ASTRAL based on arguments presented by Simmons et al. (2019).

**Figure 1.**
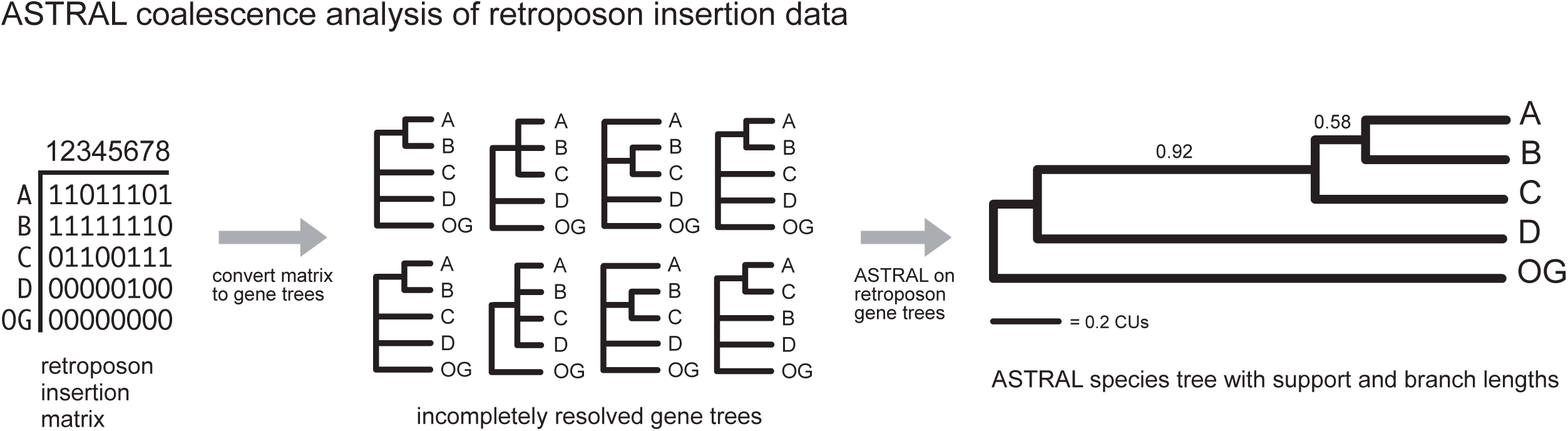
The methodology employed here for analysis of retroposons within a multispecies coalescent context. First, convert a presence/absence matrix for retroposon insertions to incompletely resolved gene trees. Next, use ASTRAL analysis of these gene trees to estimate the species tree. For the hypothetical data set in this example, posterior probabilities are above internal branches of the species tree. Branch lengths in coalescent units (CUs) are indicated for these two branches (see scale bar). Terminal branch lengths in the tree are arbitrary and are not estimated in ASTRAL analysis.

Palaeognathae includes volant tinamous and the flightless ratites (ostrich, emu, rheas, kiwis, cassowaries). Recent analyses based on phylogenomic data sets have recovered conflicting phylogenetic hypotheses for this clade (Yonezawa et al., 2017; Cloutier et al., 2019; Sackton et al., 2019). All of these studies recovered a basal split between ostrich and other palaeognaths, but relationships among the non-ostrich palaeognaths differ in Yonezawa et al. (2017) versus the other two studies (Cloutier et al., 2019; Sackton et al., 2019). Yonezawa et al. (2017) suggested that rheas are the sister taxon to all other non-ostrich palaeognaths based on a supermatrix analysis of ∼871 kb (also see Prum et al., 2015), whereas Cloutier et al. (2019) and Sackton et al. (2019) recovered tinamous as the extant sister group to all other extant non-ostrich palaeognaths based on summary coalescent analyses with MP-EST and ASTRAL on 20850 loci. Sackton et al. (2019) attributed Yonezawa et al.’s (2017) result to the ill effects of concatenation. This assertion was corroborated by supermatrix analyses of the 20850 loci compiled by Sackton et al. (2019) that also placed rheas sister to all other non-ostrich palaeognaths. Sackton et al. (2019) and Cloutier et al. (2019) suggested that very short branches on their MP-EST species tree (Fig. 2A) provide evidence for an “empirical anomaly zone” in palaeognaths that misleads concatenation analyses. Specifically, the consecutive and very short branches of 0.3657, 0.0632, and 0.0662 CUs on their MP-EST tree are the primary evidence for asserting that the palaeognath tree resides in the anomaly zone (Sackton et al., 2019). The two shortest branches are even shorter (0.0194, 0.0532) on Cloutier et al.’s (2019) ASTRAL species tree. Consistent with this hypothesis, the most common gene tree in their analysis differs from their inferred species tree (Fig. 3A; Degnan and Rosenberg, 2006, 2009). Sackton et al. (2019) also reported that retroposon insertions from Cloutier et al. (2019) provide independent support for their preferred phylogenetic hypothesis, although the retroposons were not analyzed with ILS-aware methods in either study.

**Figure 2.**
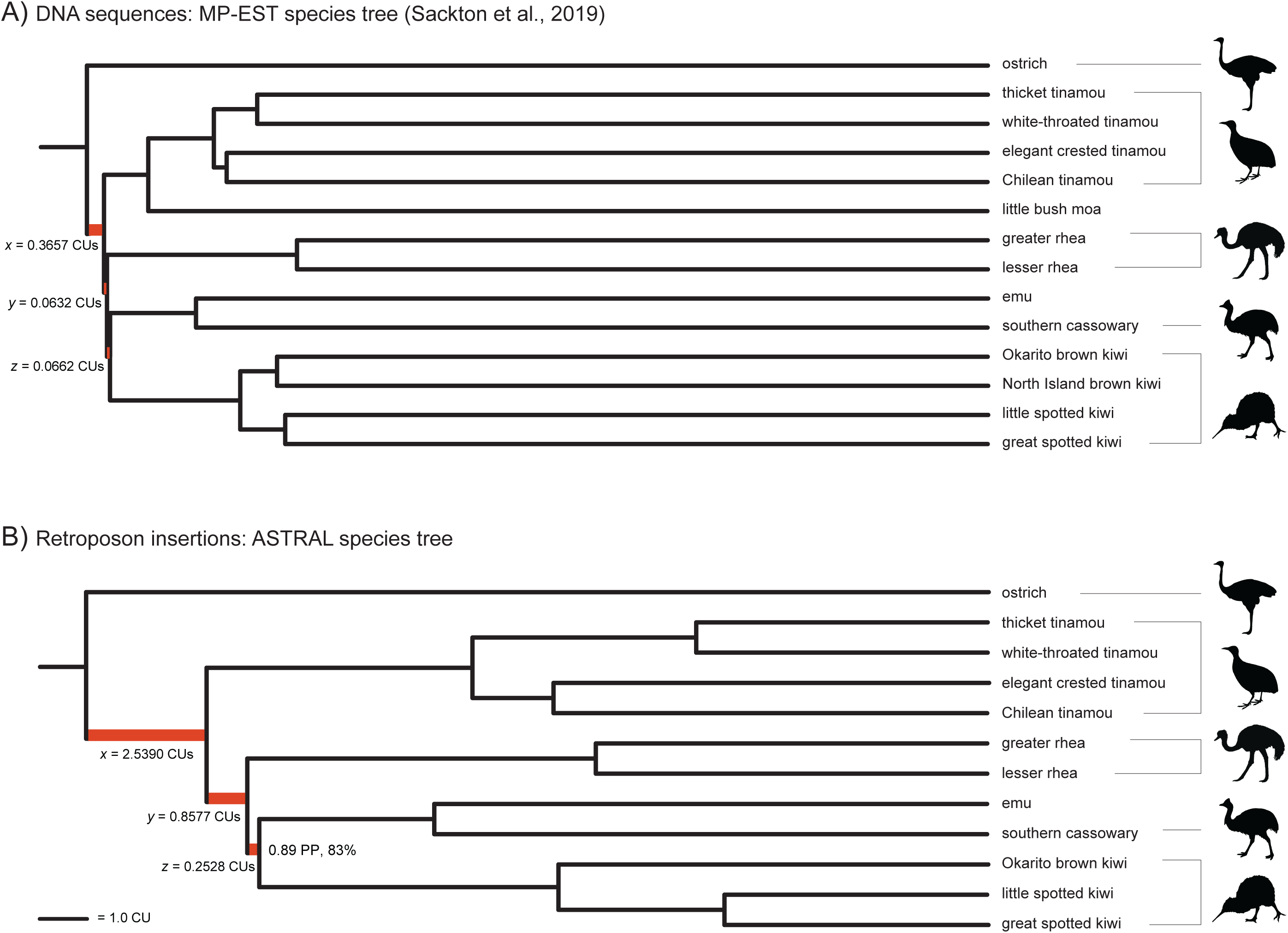
Two species tree estimates for Palaeognathae based on summary coalescence methods: (A) MP-EST species tree derived from analysis of 20850 sequence-based gene trees (Sackton et al., 2019), and (B) ASTRAL species trees derived from analysis of 4301 retroposon insertions. Note the disparity of internal branch lengths between the two trees at the base of Palaeognathae, with much longer branches for the ASTRAL tree. Scale bar in coalescent units (CUs) is identical for internal branches of both trees. The three internal branches that comprise the “empirical anomaly zone” of Sackton et al. (2019) are colored red and labeled *x*, y, and *z* as in Rosenberg (2013). Terminal branch lengths are arbitrary in both trees because MP-EST and ASTRAL do not estimate lengths of terminal branches. All clades in the MP-EST tree received 100% support for the multi-locus bootstrap (Sackton et al., 2019). All clades except one in the ASTRAL tree have local posterior probability (PP) of 1.0 and 100% support with gene-wise bootstrapping. For the one node with less than maximum support, the local PP and gene-wise bootstrap percentage are shown. Both trees are rooted with chicken (*Gallus*) outgroup (not shown), but two taxa (North Island brown kiwi, little bush moa) were not sampled in the retroposons dataset. Branch lengths for the MP-EST species tree are from the ‘TENT_mafft_MP-EST_run1.tre’ file of Sackton et al. (2019); branch lengths are similar for other MP-EST species trees posted by these authors. Silhouettes of tinamou, kiwi, and cassowary are freely available from PhylPic (http://phylopic.org/; Public Domain Mark 1.0 licence). Ostrich (credit to Matt Martyniuk [vectorized by T. Michael Keesey]; https://creativecommons.org/licenses/by-sa/3.0/) and rhea (Darren Naish [vectorized by T. Michael Keesey]; https://creativecommons.org/licenses/by/3.0/) silhouettes also were downloaded from PhyloPic.

**Figure 3.**
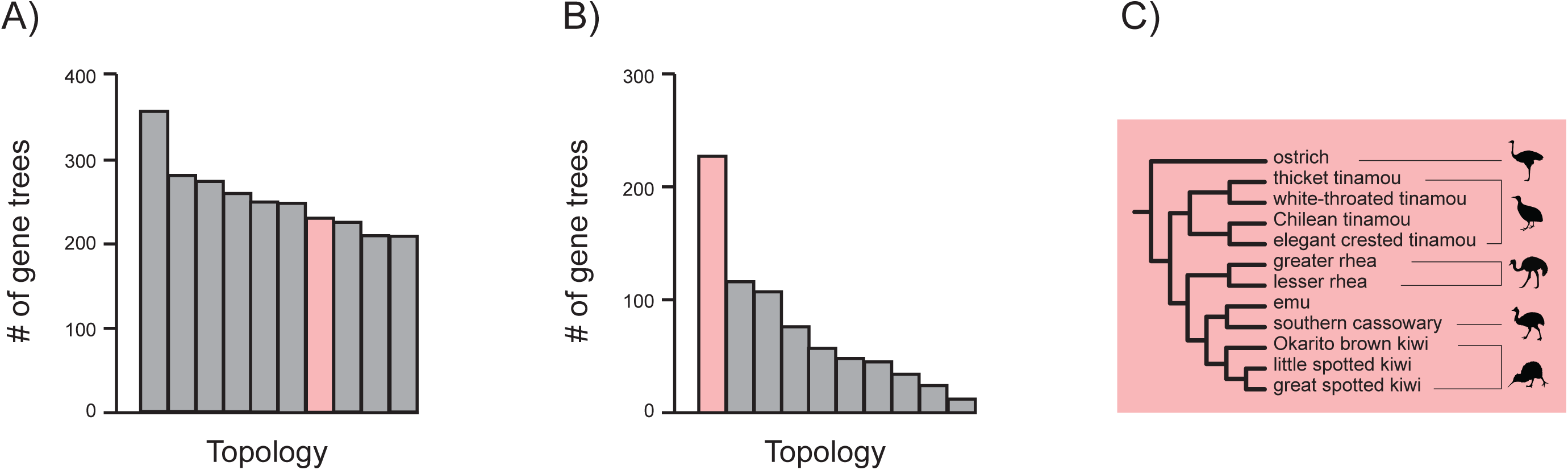
The frequencies of gene trees from Sackton et al.’s (2019) analysis of DNA sequence data from 20850 loci (A), and frequencies for 1000 gene trees simulated from the ASTRAL tree based on transposition insertions (B), with numbers of gene trees that match the species tree topology (C) highlighted in pink. In each case, bars show counts of the ten most common gene trees. For the sequence-based gene trees (modified from figure 1B of Sackton et al., [2019]), the most common gene tree does not match the species tree. The gene tree topology that matches the species tree is instead the seventh most commonly recovered topology and represents just a tiny percentage of the 20850 gene trees in this study. For gene trees simulated from the ASTRAL species tree (retroposon data), the gene tree topology that matches the species tree is by far the most common gene tree (22.7% of the gene trees that were simulated). Note that the stem Palaeognathae branch was not directly estimated on the ASTRAL retroposon tree, so we arbitrarily set this branch at 10 CUs for simulating gene trees. This branch length is a very conservative underestimate if we use the median divergences times for Aves (111.2 MY) and Palaeognathae (84.0 MY) on timetree.org and Patel et al.’s (2013) conversion of 1 CU = 0.4 MY for many vertebrates. For the long stem Palaeognathae branch (∼27.2 MY), deep coalescences due to ILS between the deeply divergent chicken lineage and various paleognath species is assumed to be rare. The topology in (C) shows taxa that were sampled for both transposons and DNA sequences; two additional species that were sampled for DNA sequences are not shown.

We re-analyzed the retroposon data set (4301 insertion characters) with ASTRAL and the resulting species tree is robustly supported (Fig. 2B). All relationships agree with the MP-EST species tree based on DNA sequence alignments (Fig. 2A; Sackton et al., 2019), but a fundamental difference is that internal branch lengths (in CUs) are much longer on the ASTRAL retroposon species tree (Fig. 2B). On average, internal branch lengths on the retroposon species tree are ~4.8X longer than the corresponding internal branches on the MP-EST species tree, including the three branches that make or break the anomaly zone (Fig. 2). First, the common ancestral branch for all palaeognaths excepting ostrich is 2.539 CUs on the retroposon tree but only 0.3657 CUs on the MP-EST tree. Second, the most striking discrepancy is for the internal branch leading to rheas, kiwis, emu, and cassowary where the ASTRAL retroposon branch length (0.8577 CUs) is ~13.6X longer than the same branch (0.0632 CUs) on the MP-EST tree constructed from DNA sequence-based gene trees. Finally, the internal branch leading to kiwis, emu, and cassowary is 0.2528 CUs on the ASTRAL tree but only 0.0662 CUs on the MP-EST tree. We used the ASTRAL retroposon species tree with its estimates of branch lengths in CUs to simulate gene trees using the program DendroPy v3.12.0 (Sukumaran and Holder, 2010) and a script from Mirarab et al. (2014b). A sample of one thousand simulated gene trees shows that the most common gene tree topology is perfectly congruent with the ASTRAL species tree and accounts for 22.7% of the simulated trees (Fig. 3B). The next most common tree was a distant 11.6% of the total. These results contrast strikingly with the distribution of gene tree topologies based on analyses of DNA sequence data, for which only ~240 of the 20850 gene trees match the MP-EST species tree (Fig. 3A; Cloutier et al., 2019; Sackton et al, 2019).

As noted above, the consecutive short branches of 0.3657, 0.0632, and 0.0662 CUs on the palaeognath MP-EST tree are the primary evidence for asserting that this species tree resides in an “empirical anomaly zone” (Sackton et al, 2019). If we view the early palaeognath radiation as a five-taxon problem (ostrich, tinamous, rheas, emu + cassowaries, and kiwis), then the three internal branches that define relationships among these lineages may each have a maximum value of 0.1934 CUs in the anomaly zone if they are coequal in length (Rosenberg and Tao, 2008; Degnan and Rosenberg, 2009). One or two of these branches may be longer in the anomaly zone, but only if the other branches compensate by being shorter (see length limits for branches *x, y*, and *z* in figure 4 of Rosenberg, 2013). In the case of a four-taxon problem (tinamous, rheas, emu + cassowaries, and kiwis), the two internal branches may each have a maximum value of 0.156 CUs if they are coequal and in the anomaly zone (Degnan and Rosenberg, 2006). If the basal branch length (*x*) is > 0.2655 then the anomaly zone vanishes for the four-taxon case (Degnan and Rosenberg, 2006). Sackton et al.’s (2019) MP-EST tree has three consecutive branches (*x, y, z*) that are consistent with an anomaly zone situation for five taxa (Rosenberg, 2013), including two branches that are < 0.1934 CUs (Fig. 2A). Similarly, Sackton et al.’s (2019) MP-EST tree is consistent with an anomaly zone situation for four taxa. However, there are no internal branches on our ASTRAL retroposon tree that are below 0.1934 CUs (Fig. 2b) for the five-taxon case or below 0.156 CUs for the four-taxon case. We used gene-wise bootstrap resampling (Simmons et al., 2019) with ASTRAL to pseudoreplicate 100 species trees with branch lengths in CUs. In every case, the branch lengths (*x, y, z*) on these species trees are not short enough to warrant an anomaly zone interpretation in either the four or five-taxon case.

**Figure 4.**
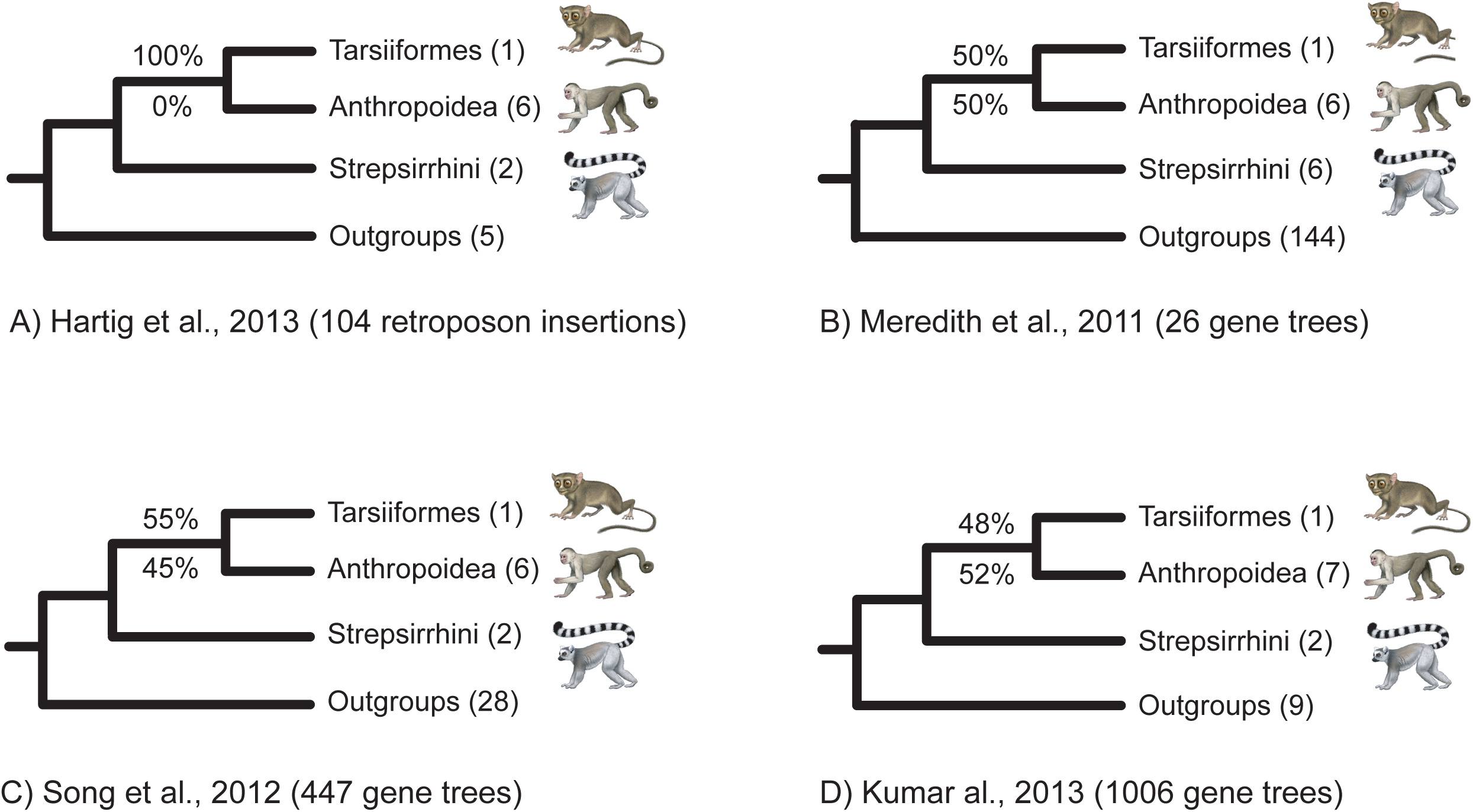
Conflict and congruence for the phylogenetic placement of Tarsiiformes (tarsiers) within Primates in recent large-scale systematic studies (A-D). Phylogenetic analyses of retroposon insertions (A) and DNA sequences (B-D) robustly position Tarsiiformes as sister to Anthropoidea. For the 104 retroposons insertions that were sampled, 100% congruence with no conflicts implies an extremely low frequency of ILS, because retroposon insertion data are thought to have low homoplasy, and no conflicts were detected at this node. DNA-based reconstructions of gene trees in multiple studies (B-D) show a very different pattern with just 48%-55% of sequence-based gene trees congruent (above internodes) and 45%-52% of these gene trees in conflict with the Tarsiiformes + Anthropoidea clade (below internodes). This suggests rampant ILS and/or gene flow where no conflicts among gene trees are expected based on the retroposon insertion data. A much simpler explanation is wholesale gene tree reconstruction error in the three sequence-based analyses (B-D). For each study, the number of gene trees reconstructed is shown, and the number of species sampled for each group is in parentheses to the right of each taxonomic name. ‘Outgroups’ refers to all species in an analysis that are not primates. Illustrations of primates are by C. Buell.

The internal branch lengths in the sequence-based species tree (Fig. 2A) and the retroposon-based species tree (Fig. 2B) tell two very different stories about the early diversification of palaeognaths. For example, 18 retroposon insertions show uniform support, with no conflicts, for the clade that includes all palaeognaths except the ostrich (Cloutier et al., 2019), suggesting limited ILS. By contrast, sequence-based gene trees show extensive conflict at this node (69% for conserved non-exonic elements, 39% for UCEs, 35% for introns). By contrast, absolutely no derived retroposon insertions were found in ostrich that conflict with this clade – i.e., 0% conflict (Cloutier et al., 2019). We contend that the much longer internal branch lengths on the retroposon species tree for this clade and others are more accurate than the much shorter branch lengths on Sackton et al.’s (2019) MP-EST species trees. This is because gene tree reconstruction error is negligible for retroposons (Doronina et al., 2019), but not for sequence-based gene trees (Huang and Knowles, 2009; Huang et al., 2010; Springer and Gatesy, 2014, 2016, 2018a, b; Bayzid et al., 2015; Mirarab and Warnow, 2015; Simmons and Gatesy, 2015; Sayyari et al., 2016; Gatesy et al., 2017 Sayyari and Mirarab, 2017; Scornavacca and Galtier, 2017; Molloy and Warnow, 2018). Cloutier et al. (2019) noted that an “important and related question is whether we can confidently detect an empirical anomaly zone given that the same short internal branches expected to produce anomalous gene trees also increase the likelihood of gene tree estimation error due to the short time interval for informative substitutions to accumulate in individual loci of finite length [Huang and Knowles, 2009].” Ironically, the retroposon data collected by these authors provide a definitive answer to this question when analyzed within the multispecies coalescent with the program ASTRAL (Fig. 2B).

Previous studies repeatedly have shown that branch lengths (in CUs) on coalescence-based species trees are too short because of gene tree reconstruction error, which mimics conflict due to ILS (Mirarab et al., 2014a,b; Gatesy and Springer, 2014; Springer and Gatesy, 2014, 2016, 2018a,b; Bayzid et al., 2015; Roch and Warnow, 2015). Sequence-based gene trees and retroposon-based ‘gene trees’ for the phylogenetic placement of tarsiers within Primates provide a stellar example of the gene-tree reconstruction error problem. In this case, large-scale phylogenetic and phylogenomic analyses of DNA sequences have consistently recovered a sister-group relationship between Tarsiiformes (tarsiers) and Anthropoidea (monkeys + apes) to the exclusion of Strepsirrhini (lemurs + lorises) (Fig. 4B-D; Meredith et al., 2011; Song et al., 2012; Kumar et al., 2013). A sister-group relationship between Tarsiiformes and Anthropoidea is also supported by 104 retroposon insertions without any conflict among the retroposon characters (Fig. 4A; Hartig et al., 2013). The absence of any conflicting retroposons suggests that ILS is negligible for this node that is deep in the primate tree (Springer and Gatesy, 2018b). By contrast, DNA-based reconstructions of individual gene trees exhibit poor support for Tarsiiformes + Anthropoidea, with 45%-52% of gene trees conflicting with this clade (Fig. 4).

More generally, we argue that much of the gene tree heterogeneity in sequence-based analyses at deep divergences is unrelated to ILS and instead results from long branch misplacement, model misspecification, lack of phylogenetic signal, missing data, homology problems, and poor tree searches (Gatesy and Springer, 2013, 2014; Patel et al., 2013; Springer et al., 2014, 2016, 2017 2018a-c; Gatesy et al., 2017, 2018). Such gene tree reconstruction errors artificially inflate conflicts and are known to result in highly stunted internal branches on MP-EST species trees (Gatesy and Springer, 2014; Mirarab et al., 2014a,b; Springer and Gatesy, 2014, 2016, 2018b; Bayzid et al., 2015; Roch and Warnow, 2015). In the case of palaeognaths, the much longer internal branches on our retroposon species tree therefore call into question the existence of an “empirical anomaly zone” for Palaeognathae.

Cloutier et al. (2019) used simulations of gene trees from their MP-EST species tree with tiny internal branches (Fig. 2A) to “assess what proportion of total gene tree heterogeneity is likely attributable to coalescent processes rather than to gene tree estimation error” and found that 70%-90% of the gene tree conflict is compatible with ILS. However, previous work has shown that this simulation procedure is flawed because of gene tree reconstruction error that stunts branches on sequence-based MP-EST species trees (Springer and Gatesy, 2014, 2016, 2018a,b). By contrast, retroposon ‘gene trees’ can be more reliably reconstructed due to low levels of homoplasy at single genomic sites. In accordance with this expectation, the palaeognath species tree based on retroposons has ~4.8X longer internal branch lengths than the species tree that was reconstructed from sequence-based gene trees (Cloutier et al., 2019; Sackton et al., 2019). Furthermore, branch lengths on the retroposon species tree for palaeognaths are inconsistent with the “empirical anomaly zone” hypothesis (Figs. 2B, 3C). To our knowledge, Linkem et al.’s (2016) analysis of skink phylogeny is the only other study that reports a putative empirical example of the anomaly zone. However, Linkem et al.’s (2016) argument was partly based on MP-EST branch lengths, which are too short when there is sequence-based gene tree error that mimics ILS (Mirarab et al., 2014a,b; Springer et al., 2014, 2016; Bayzid et al., 2015; Roch and Warnow, 2015). In view of our results, which demonstrate striking branch length differences between sequence-based and retroposon-based species trees, we suggest that the former will have a high false positive rate for the discovery of putative cases of anomaly zone branch lengths. We suggest that retroposon insertions, analyzed in a multispecies coalescent framework, provide a more reliable approach for the identification of candidate anomaly zone topologies.

Summary coalescence methods explicitly address the problem of incomplete lineage sorting, which can negatively impact the performance of concatenation approaches for species tree reconstruction. However, these methods assume no intralocus recombination, neutral evolution, and no gene tree reconstruction error. In challenging areas of treespace such as the anomaly zone (Degnan and Rosenberg, 2006; Rosenberg, 2013), summary coalescence methods are not guaranteed to provide reliable evolutionary inferences if these assumptions are violated. This problem is especially acute for branch lengths estimates, which are directly derived from the level of gene tree conflict (Liu et al., 2010; Sayyari and Mirarab, 2016) and will therefore be stunted when gene tree error mimics ILS (Mirarab et al., 2014a; Springer and Gatesy, 2014). We contend that the key assumptions of summary coalescence methods are in much better agreement with retroposon insertions than with sequence-based gene trees. In particular, branch lengths in CUs (Fig. 2) are expected to be more accurate when the underlying data have lower levels of homoplasy and internal recombination, as is the case for retroposons (Fig. 4). Coalescence analysis of retroposon insertions (Fig. 1) therefore provides a promising alternative to the status quo of estimating species trees based on sequence-based gene trees, especially at deep divergences where homoplasy is prevalent for characters with just four states (i.e., DNA sequences). Indeed, the application of summary coalescence methods to retroposon data may effectively liberate summary coalescence methods from their greatest hindrance -gene tree reconstruction error (Huang et al., 2010). The procurement of retroposon data sets and the development of novel multispecies coalescent methods for the analysis of these data sets should be a priority for future phylogenomic studies.

## Acknowledgments

C. Buell provided images of mammals. This research was funded by NSF DEB-1457735.

